# Identifying cis-mediators for trans-eQTLs across many human tissues using genomic mediation analysis

**DOI:** 10.1101/078683

**Authors:** Fan Yang, Jiebiao Wang, the GTEx consortium, Brandon L. Pierce, Lin S. Chen

## Abstract

The impact of inherited genetic variation on gene expression in humans is well-established. The majority of known expression quantitative trait loci (eQTLs) impact expression of local genes (cis-eQTLs). More research is needed to identify effects of genetic variation on distant genes (trans-eQTLs) and understand their biological mechanisms. One common trans-eQTLs mechanism is “mediation” by a local (cis) transcript. Thus, mediation analysis can be applied to genome-wide SNP and expression data in order to identify transcripts that are “cis-mediators” of trans-eQTLs, including those “cis-hubs” involved in regulation of many trans-genes. Identifying such mediators helps us understand regulatory networks and suggests biological mechanisms underlying trans-eQTLs, both of which are relevant for understanding susceptibility to complex diseases. The multi-tissue expression data from the Genotype-Tissue Expression (GTEx) program provides a unique opportunity to study cis-mediation across human tissue types. However, the presence of complex hidden confounding effects in biological systems can make mediation analyses challenging and prone to confounding bias, particularly when conducted among diverse samples. To address this problem, we propose a new method: Genomic Mediation analysis with Adaptive Confounding adjustment (GMAC). It enables the search of a very large pool of variables, and adaptively selects potential confounding variables for each mediation test. Analyses of simulated data and GTEx data demonstrate that the adaptive selection of confounders by GMAC improves the power and precision of mediation analysis. Application of GMAC to GTEx data provides new insights into the observed patterns of cis-hubs and trans-eQTL regulation across tissue types.

## INTRODUCTION

Recent studies have demonstrated that many expression quantitative trait loci (eQTLs) that affect expression of local transcripts (cis-eQTLs) also affect the expression of distant genes (trans-eQTLs) (Battle et al. 2014; Pierce et al. 2014). This observation suggests the effects of trans-eQTLs are “mediated” by the local (cis-) gene transcripts near the eQTLs (Fehrmann et al. 2011; Pierce et al. 2014). In other words, some cis-eQTLs are also trans-eQTLs because the variation in the expression of the cis-gene effects the expression of a trans-gene or genes. In the simplest scenario, a cis-eQTL effects expression of a nearby gene that is a transcription factor, which then regulates the transcription of a distant gene; thus, the transcription factor “mediates” the effect of the eQTL on the distant gene. By studying eQTLs that have both the cis- and trans-effects, one may identify the cis-genes that mediate the effect of a trans-eQTL on expression of a distant gene, including “cis-hub” genes that regulate the expression of many trans-genes (Chen et al. 2007; Stranger et al. 2012). Studying mediation (causation) moves beyond the analysis of cis- and trans- associations (correlation). Prior studies have applied mediation tests to genome-wide SNP and expression data (from blood cells) to identify transcripts that are cis-mediators of the effects of trans-eQTLs (Chen et al. 2007; Battle et al. 2014; Pierce et al. 2014). Characterizing these regulatory relationships will allow us to better understand regulatory networks and their roles in complex diseases (Veyrieras et al. 2008), as it is well-known that SNPs influencing human traits tend to be eQTLs (Nicolae et al. 2010). Analyses of cis-mediation will also provide us with a better understanding of the biological mechanisms underlying trans-eQTLs (Westra et al. 2013).

The expression levels of a given gene can vary substantially across human cell types, and the regulatory relationships between SNPs and gene expression levels may also depend on cell type (Torres et al. 2014; Wang et al. 2016). To date, most large-scale eQTL studies have been conducted using RNA extracted from peripheral blood cells, which are mixtures of different cell types and may not be informative for gene regulation in other human tissues. In order to study gene expression and regulation in a variety of human tissues, the National Institutes of Health common-fund GTEx (Genotype-Tissue Expression) project has collected expression data on 44 tissue types from hundreds of post-mortem donors (Lonsdale et al. 2013; Ardlie et al. 2015). This rich transcriptome data, coupled with data on inherited genetic variation, provides an unprecedented opportunity to study gene expression and regulation patterns from both cross-tissue and tissue-specific perspectives.

One major challenge in mediation analysis is the presence of unmeasured or unknown variables that affect both the mediator (i.e., cis-gene) and outcome (i.e., trans-gene) variables. This presence of such a variable is known as “mediator-outcome confounding,” and in such a scenario, estimates obtained from mediation analysis can be biased (Robins and Greenland 1992; Pearl 2001; Cole and Hernan 2002). In other words, in the presence of an unmeasured confounding variable(s), the association between the two cis- and trans-genes will be a biased estimate of the causal relationship between the two genes, and estimates obtained from mediation analysis will be biased. It is well recognized that measures of transcriptional variation can be affected by genetic, environmental, demographic, technical, as well as biological factors. The presence of unmeasured or unknown confounding effects may induce inflated rates of false detection of mediation relationships or jeopardize the power to detect real mediation, if those confounding effects are not well accounted for. Given that eQTL analyses are conducted in the context of complex biological systems, there are a wide array of biological variables that could potentially confound the mediator-outcome association and bias mediation estimates, a problem that may be exacerbated by the diversity of GTEx participants, with respect to ethnicity, age, and cause of death. Given these challenges, it is desirable to have methods that consider a large pool of potential confounding variables.

To adjust for unmeasured or unknown confounding effects in genomics studies, existing literature focused on the construction of sets of “hidden” variables that capture a substantial amount of the variation in a large set of variables (Price et al. 2006; Leek and Storey 2007; Stegle et al. 2012). A commonality of those approaches is that they model the effects of hidden confounding factors and summarize those effects into a set of constructed variables, sort those variables decreasingly by their estimated impacts, and adjust the top ones as a set of covariates to eliminate major confounding effects in the subsequent analysis. For example, in GTEx eQTL analyses (Ardlie et al. 2015) (cite Jo et al., GTEx companion paper, unpublished) the top Probabilistic Estimation of Expression Residuals (PEER) factors were estimated for each tissue type and up to 35 factors were adjusted. One aspect that is largely ignored is that not all potential gene pairs (or pairs of regulator and regulated genes) are affected by the same set of hidden confounders. There are likely thousands of cis-mediated trans-eQTLs in the human genome, i.e., trios consisting of a genetic variant, a cis-gene transcript, and a trans-gene transcript in a specific tissue type. However, for each trio, mediator-outcome confounding will be present only when a hidden variable is causally related to the regulator and regulated genes. By this criterion, the potential confounder set varies by different trios. Adjusting a universal set of variables for all mediation trios is not only inefficient but also may limit our ability to consider a larger pool of potential confounding variables in genomic mediation analyses.

We propose to adaptively select the variables to adjust for each trio given a large set of constructed or directly measured potential confounding variables. This strategy supplements existing confounding adjustment approaches that focus on the construction of variables for capturing confounding effects, and enlarges the pool of variables to be considered. Additionally, by leveraging the cis genetic variant as an “instrumental variable,” we are able to select the variables capturing confounding effects rather than variables only correlated with cis- and trans-genes. We further propose a mediation test with non-parametric *p*-value calculation, adjusting for the adaptively selected sets of confounders. We term the proposed algorithm Genomic Mediation analysis with Adaptive Confounding adjustment (GMAC). The GMAC algorithm improves the efficiency and precision of confounding adjustment and the subsequent genomic mediation analyses. We applied GMAC to each of the 44 tissue types of GTEx data in order to study the trans-regulatory mechanism in human tissues. Our algorithm identifies genes that mediate trans-eQTLs in multiple tissues, as well as “cis-hubs” that mediate the effects of a trans-eQTL on multiple genes.

## RESULTS

### GMAC improves power and precision of analysis of GTEx data

We performed genomic mediation analysis with data from each tissue type in GTEx. Taking the tissue Adipose Subcutaneous as an example, there are 298 samples for this tissue type and gene-level expression measures for 27,182 unique transcripts are available after quality control. Consider a candidate mediation trio consisting of a gene transcript *i* (*C_i_*), its cis-associated genetic locus (*L_i_*), and another gene transcript *j* (*T_j_*) in trans-association with the locus. The goal is to test for mediation of the effect of the genetic locus on the trans-gene by the cis-gene (see Figure 1). We focused on only the trios (*L_i_*, *C_i_*, *T_j_*) in the genome showing both cis- and trans-eQTL associations, i.e., *L_i_ → C_i_* and *L_i_ → T_j_*. Because associations are necessary but not sufficient conditions for inferring mediation, by considering only the trios with cis- and trans-associations, we effectively reduced the search space to a promising pool of candidate mediation trios and alleviated the multiple testing burdens. We detected and selected a lead cis-eQTL for 8,500 of these transcripts, corresponding to 8,216 unique cis-eSNPs for subsequent analysis (see Methods). We applied Matrix eQTL (Shabalin 2012) to the 8,216 SNPs and the 27,182 gene expression levels to calculate the pair-wise trans-associations. At the *p*-value cutoff of 10^−5^, there were 3,169 significant pairs of SNP and trans-gene transcripts. Since some cis-eSNPs were the lead cis-eSNPs for multiple local gene transcripts, those significant SNP and trans-gene pairs entailed a total of 3,332 trios (i.e., SNP-cis-trans) for this tissue type. We applied GMAC (see Methods and see Figure 2 for a graphical illustration of the main steps of the GMAC) to the 3,332 trios in this tissue type to test for mediation, and obtained the mediation *p*-values for those trios. Since different tissue types have different sample sizes in GTEx and in addition to cross-tissue confounders there are many tissue-specific confounding effects, we constructed Principal Components (PCs) from the expression data of each tissue type as potential confounders (Figure 2B). The number of PCs for each tissue type is equal to the tissue sample size minus 1. We analyzed trios for mediation in a similar fashion for all other GTEx tissue types.

**Figure 1.**
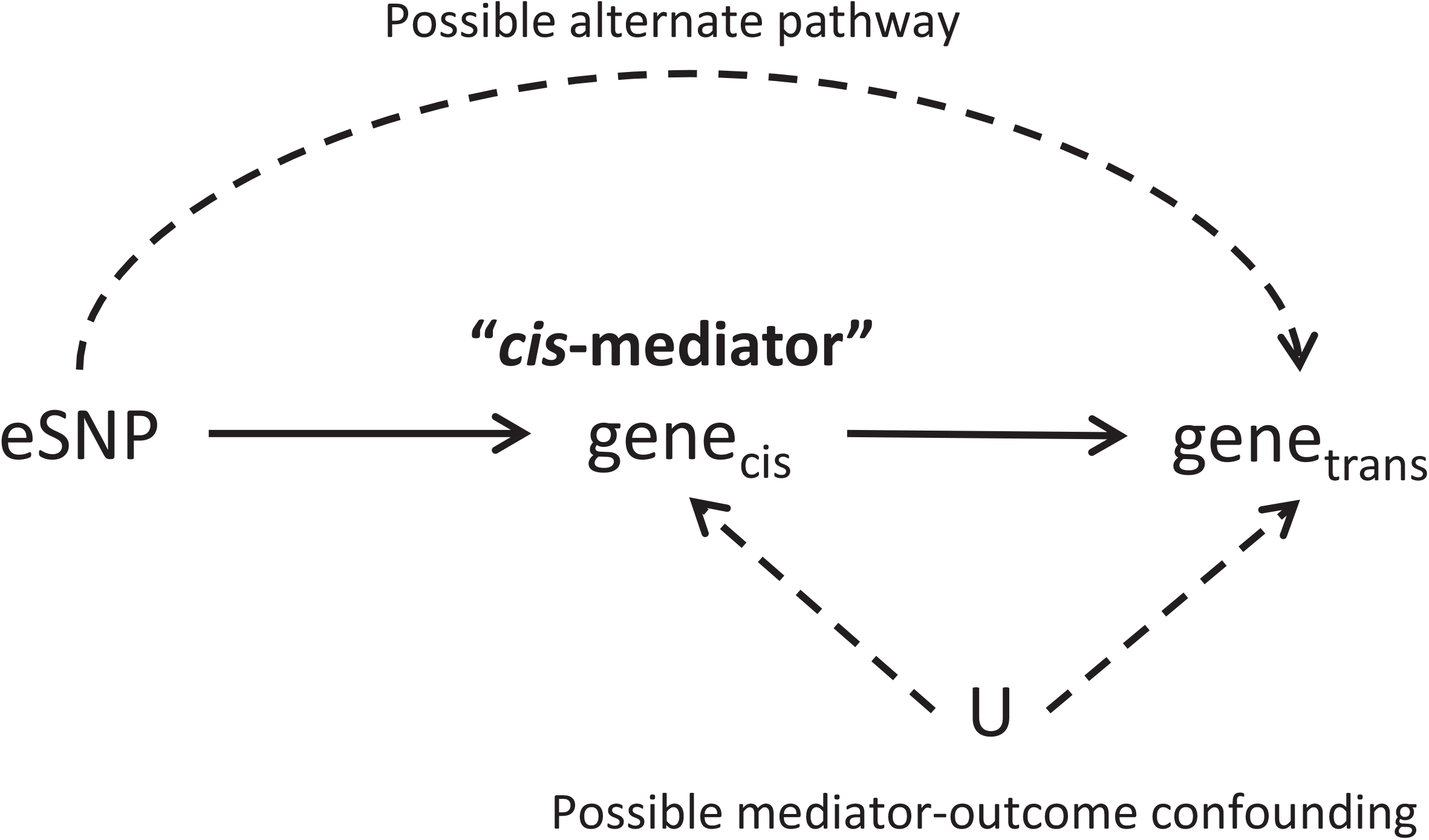
Causal diagram demonstrating mediation and “mediator-outcome confounding.” Here the variable set “**U**” represents a set of unmeasured or unknown variables that may show confounding effects in the mediation analysis. Mediation analysis can detect mediation of the effect of the eSNP on the trans- gene by the cis-gene, assuming mediator-outcome confounding is absent or adjusted for in the analysis. Mediation will not be detected if the effect of the eSNP on the trans-gene is through some alternative pathway that does not involve the cis-gene.

**Figure 2.**
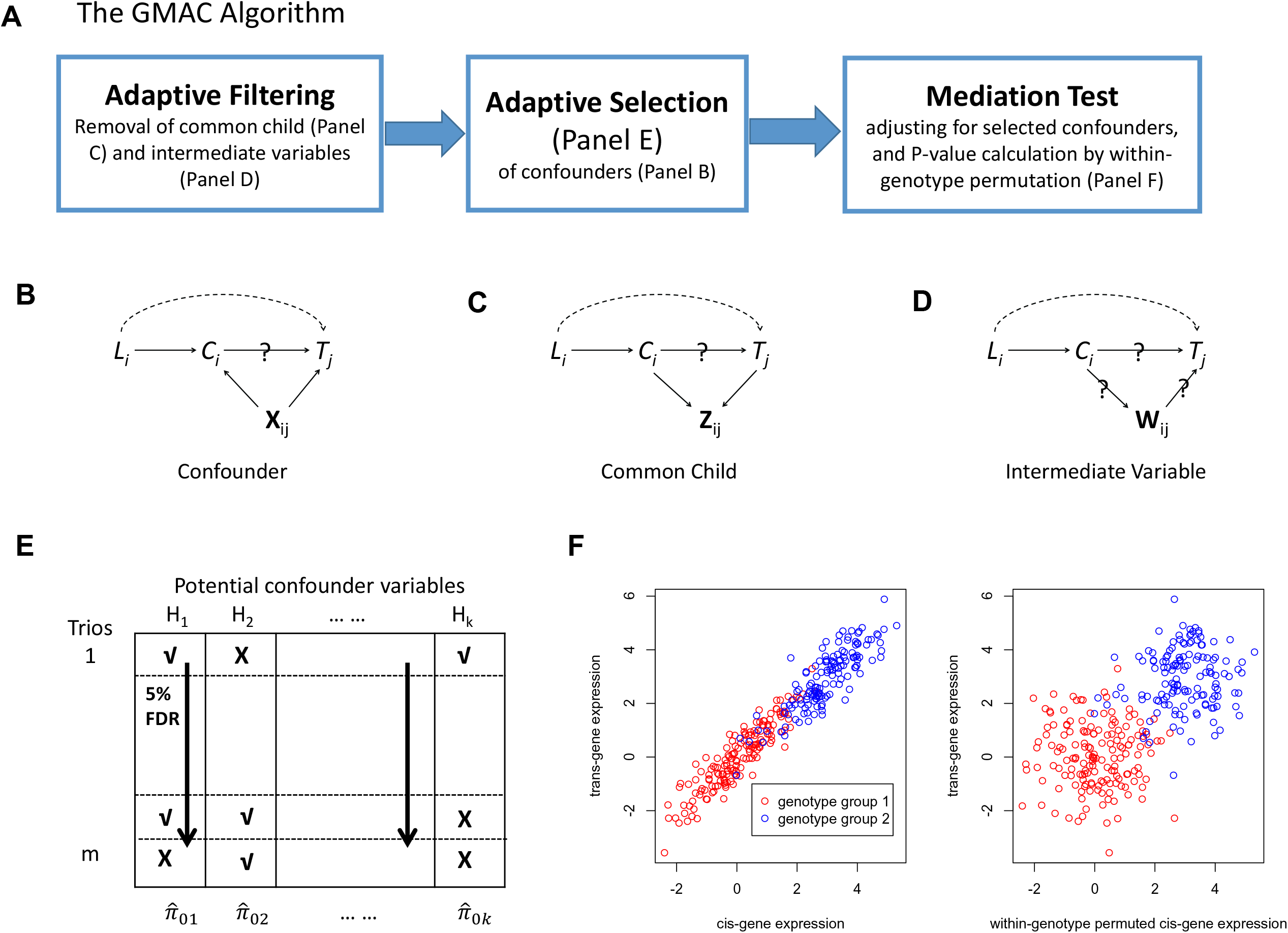
Graphical illustrations of GMAC and its main idea. (A) A summary of the GMAC algorithm; (B) a mediation relationship among an eQTL, *L_i_*, its cis-gene transcript, *C_i_*, and a trans-gene transcript, *T_j_*, with confounders, **X_ij_**, allowing *L_i_* to affect *T_j_* via a pathway independent of *C_i_*; (C) a mediation trio where *C_i_*, and *T_j_*, have common child variable(s), **Z_ij_**; (D) a mediation trio where *C_i_*, affects *T_j_*, through intermediate variable(s), **W_ij_**. (E) The adaptive confounder selection procedure: Based on the *p*-value matrix for the association of each potential confounder variable to at least one of the cis- or the trans-gene transcript, we apply a stratified FDR approach by considering the *p*-values for each potential confounder (each column) as a stratum, with the significant ones indicated by a check mark (√). When conducting the mediation test for each trio, we only adjust for the significant confounding variables (the ones with with in each row). (F) A mediation trio *L_i_* → *c_i_* → *T_j_* (left) and a trio under the null with both cis-linkage and trans-linkage but no mediation (right). Within-genotype permutation of the cis-gene expression levels maintains the cis- and trans-linkage (different mean levels) while breaks the potential correlation between the cis- and trans-expression levels within each genotype group. Note that **X_ij_**, **Z_ij_**, **W_ij_** may vary by trios and are all subsets of **H**. We assume that either **X_ij_** or a combination of variables in **X_ij_** would capture the variation of the unmeasured confounder **U** in Figure 1.

At the 5% false discovery rate (FDR) (Storey and Tibshirani 2003) level, we identified 6,145 instances of significant mediation out of 64,824 trios tested in the 44 tissue types. These trios represent potential examples of cis-mediation of trans-eQTLs within a specific tissue. Table 1 lists the number of significant mediation trios at 5% FDR and the number of trios with suggestive mediation (*p*-value < 0.05), as well as the total number of trios with significant cis- and trans-associations for all tissue types. The number of confounders selected for each mediation test ranged from 0 to 22 across all tissue types, with a mean of 7.695 and a median of 8. The median number of confounders selected for each tissue type ranged from 3 to 12, while the pool of variables (PCs) from which we selected confounders from ranged from 69 to 360. Supplemental Table S1 presents the descriptive statistics for the number of selected confounders for all the trios in each tissue type. It is clear that with GMAC, on average we adjust an efficient number while considering a large pool of confounding variables in the mediation tests, and that may improve the power and accuracy of the analyses.

**Table 1.** A description of GTEx tissue types and the number of significant instances of mediation (i.e., SNP-cis-trans trios) identified by GMAC (Genomic Mediation Analysis with adaptive Confounder adjustment).

| Tissue name | Tissue sample size | # Trios tested | # Trios with suggestive mediation ($p$-value <0.05) | # Trios significant at 5% FDR |
| --- | --- | --- | --- | --- |
| Muscle Skeletal | 361 | 2387 | 496 | 264 |
| Whole Blood | 338 | 2274 | 508 | 281 |
| Skin Sun Exposed Lower leg | 302 | 3273 | 629 | 330 |
| Adipose Subcutaneous | 298 | 3332 | 640 | 325 |
| Artery Tibial | 285 | 2699 | 527 | 281 |
| Lung | 278 | 2762 | 543 | 323 |
| Thyroid | 278 | 3894 | 696 | 376 |
| Cells Transformed fibroblasts | 272 | 3000 | 642 | 340 |
| Nerve Tibial | 256 | 3812 | 677 | 326 |
| Esophagus Mucosa | 241 | 2640 | 465 | 242 |
| Esophagus Muscularis | 218 | 2431 | 447 | 230 |
| Artery Aorta | 197 | 2009 | 368 | 186 |
| Skin Not Sun Exposed Suprapubic | 196 | 1961 | 365 | 177 |
| Heart Left Ventricle | 190 | 1290 | 242 | 115 |
| Adipose Visceral Omentum | 185 | 1410 | 257 | 125 |
| Breast Mammary Tissue | 183 | 1422 | 254 | 126 |
| Stomach | 170 | 1153 | 235 | 107 |
| Colon Transverse | 169 | 1585 | 309 | 161 |
| Heart Atrial Appendage | 159 | 1221 | 243 | 103 |
| Testis | 157 | 3896 | 607 | 267 |
| Pancreas | 149 | 1270 | 208 | 102 |
| Esophagus Gastroesophageal Junction | 127 | 857 | 148 | 74 |
| Adrenal Gland | 126 | 981 | 185 | 94 |
| Colon Sigmoid | 124 | 968 | 220 | 108 |
| Artery Coronary | 118 | 874 | 191 | 95 |
| Cells EBV-transformed lymphocytes | 114 | 856 | 152 | 78 |
| Brain Cerebellum | 103 | 1295 | 187 | 84 |
| Brain Caudate basal ganglia | 100 | 763 | 139 | 61 |
| Liver | 97 | 496 | 87 | 41 |
| Brain Cortex | 96 | 754 | 134 | 45 |
| Brain Nucleus accumbens basal ganglia | 93 | 592 | 97 | 43 |
| Brain Frontal Cortex BA9 | 92 | 595 | 102 | 52 |
| Brain Cerebellar Hemisphere | 89 | 1072 | 222 | 116 |
| Spleen | 89 | 825 | 157 | 68 |
| Pituitary | 87 | 732 | 132 | 61 |
| Prostate | 87 | 474 | 101 | 54 |
| Ovary | 85 | 469 | 95 | 43 |
| Brain Putamen basal ganglia | 82 | 481 | 94 | 35 |
| Brain Hippocampus | 81 | 343 | 93 | 47 |
| Brain Hypothalamus | 81 | 342 | 74 | 41 |
| Vagina | 79 | 248 | 58 | 25 |
| Small Intestine Terminal Ileum | 77 | 434 | 82 | 39 |
| Brain Anterior cingulate cortex BA24 | 72 | 365 | 81 | 29 |
| Uterus | 70 | 287 | 62 | 25 |

Again taking the tissue Adipose Subcutaneous as an illustration, in Figure 3 we plotted the negative log base 10 of the mediation *p*-values versus the percentages of reduction in trans-effects after adjusting for a potential cis-mediator, based on mediation tests without adjusting for hidden confounders (Figure 3A) and mediation tests by GMAC considering all PCs as potential confounders (Figure 3B). The percentage of reduction in trans-effects is calculated by 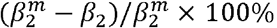, where 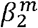 is the marginal trans-effect of the eQTL on the trans-gene expression levels, and *β*_2_ is the trans-effect after adjusting for cis-mediation. For trios representing true cis-mediation, we expect the trans-effects to be substantially reduced after adjusting for the mediator. That is, we expect the trios with very significant mediation *p*-values to have positive % reduction in the trans-effect. In Figure 3A, we observed many trios with significant mediation *p*-values, but for a substantial number of these trios, the percentages of reduction in trans-effects are negative. At the mediation *p*-value threshold of 0.05, 1577 out of 3332 trios were significant; however, 712 trios (712/3332 = 21.3%) have negative % reduction in trans-effects. This contradicting result is expected in the presence of unadjusted confounders, and many of these trios may be false positives. Thus, mediation analyses of GTEx data without adjusting for hidden confounding effects will lead to many spurious findings.

**Figure 3.**
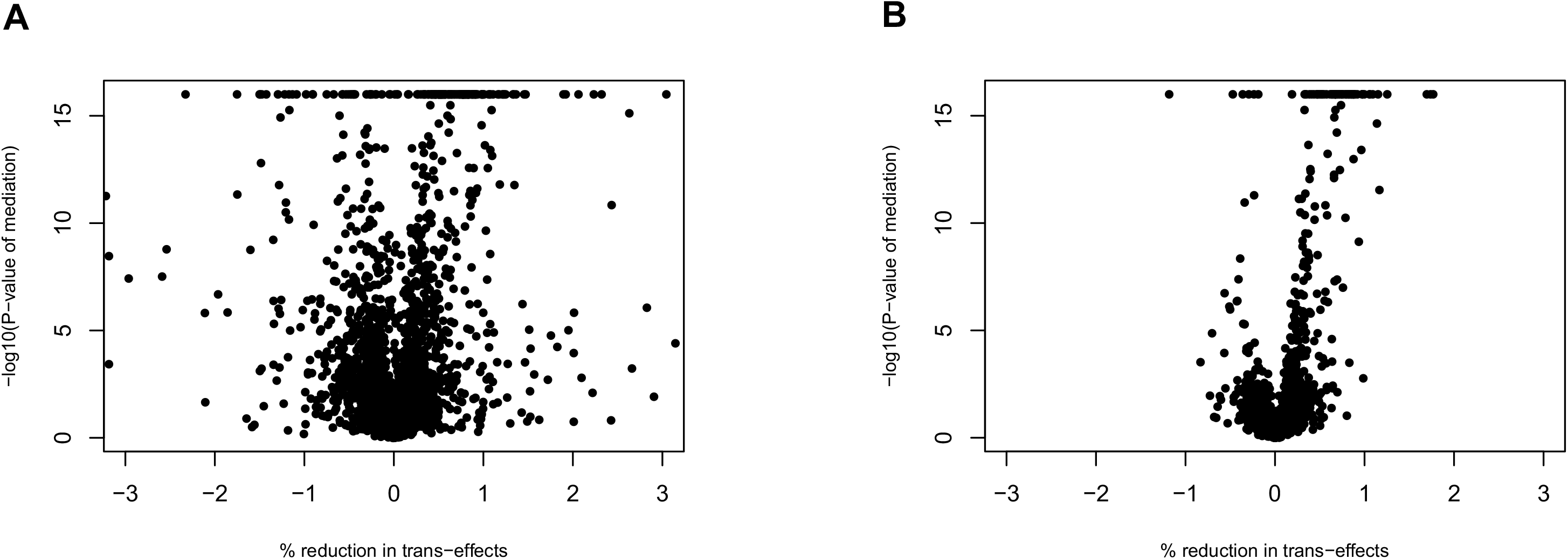
Plots of negative log base 10 of mediation *p*-values versus the percentages of reduction in trans-effects after accounting for cis-mediation. The *p*-values are calculated based on (A) mediation tests without adjusting for hidden confounders (B) mediation tests by GMAC considering all PCs as potential confounders. P-values are truncated at 10^−16^. The plots are based on the results from the Adipose Subcutaneous tissue. The percentage of reduction in trans-effects is calculated by (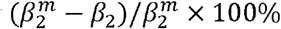), where 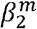 is the marginal trans-effect of the eQTL on the trans-gene expression levels, and *β*_2_ is the trans-effect after adjusted for a potential cis-mediator and other covariates. For trios with true cis- mediations, the marginal trans-effects are non-zero, and after adjusting for the true cis-mediators we expect the adjusted trans-effects *β*_2_ to be substantially reduced. That is, we expect the trios with very significant mediation *p*-values to have positive % reduction in trans-effects. For results based on no adjustment of hidden confounders (A), we observed many trios with significant mediation *p*-values but the percentages of reduction in trans-effects are negative. At the 0.05 *p*-value threshold, 712 (21.4%) and 188 (5.6%) out of 3332 trios have *p*-values below the threshold and % reduction in trans-effects being negative in Figure 3A and 3B, respectively.

In addition to our main analysis based on GMAC (adaptively selecting confounders from all expression PCs), we also conducted mediation tests adjusting for only the 35 PEER factors used in the GTEx eQTL analyses (cite Jo et al., GTEx companion paper, unpublished). At the 5% FDR level, 3,356 out of 64,824 trios from all tissue types were significant. Using GMAC adjusting for adaptively selected PEER factors, 5,131 trios were significant at the 5% FDR level. The comparison of adjusting for all (up to 35) PEER factors versus GMAC (considering a larger pool of potential confounders with up to 360 PCs) demonstrates that adaptive selection enables more efficient adjustment of confounding effects with a much fewer number of selected confounding variables (Supplemental Table S1) and improves power to detect mediation. Furthermore, using GMAC to adaptively select confounders from all PCs identifies 6,145 significant trios, suggesting an increase in power. It can also be seen that all the three methods, 1) GMAC with adaptively-selected PCs, 2) GMAC with adaptively-selected PEER factors, and 3) adjusting for all PEER factors, would yield reasonable mediation estimates (i.e., percentages of reduction in trans-effects versus mediation *p*-values), as compared to no confounder adjustment (see Supplemental Fig S1). In conclusion, motivated by the fact that the potential confounder set may vary by different trios, GMAC adaptively adjusts for only the variable that are causally related to both cis- and trans-genes and may show confounding effects in the mediation analysis of each trio (Figure 2B). Comparing with adjusting for a universal set of (top) variables for all mediation trios, GMAC considers a larger pool of potential confounding variables in genomic mediation analyses and enjoys improved power while controlling for false positives.

The majority of the cis-mediators and trans target genes observed among our trios showing mediation have high mappability scores (Supplemental Fig S2). However, non-uniquely mapping reads can result in false positive eQTLs, so we consider the mappability of each gene as a quality control filter for studying specific examples of cis-mediation (see Methods). Examining the mappability for genes involved in cis-mediation, we observed that cis-genes showing evidence of cis-mediation for multiple trans genes were enriched for cis-genes with low mappability scores (Supplemental Fig S2). Similarly, genes showing evidence of cis-mediation across many different tissue types were also enriched for genes showing low mappability scores (Supplemental Fig S2). This finding demonstrates that transcripts that do not uniquely map to the genome are an important source of false positives when conducting genomic mediation analysis. More specifically, we find that analyzing low-mappability genes can lead to the identification of spurious cis-hubs and cross-tissue cis-mediators.

We attempted to identify “cis-hubs” with high mappability in the GTEx data, defined as a transcript that appears to mediate the effect of a nearby eSNP on expression of multiple distant (i.e., trans) gene transcripts. Restricting our analysis to cis and trans genes with mappability >0.95, we observed 685 cis-genes with at least two trans targets (considering all tissues), representing 21% of the 3,168 cis-genes observed among the trios with a mediation *p*-value <0.05 (Table 2). In addition, we attempted to identify cis-genes that have at least one trans target in multiple tissues. Restricting to high mappability genes, we observed 531 cis-genes with trans targets in more than one tissue, representing 17% of the 3,168 cis-genes observed among the trios with a mediation *p*-value <0.05 (Table 2). We observed only six examples of cis-genes that had the same trans targets in multiple tissues. In other words, vast majority of cis-hubs observed were of two distinct types: 1) those that mediated the effect of a trans-eQTL on multiple trans-genes within a single tissue type, and 2) those that were mediators in multiple tissues, but with unique trans targets in each tissue type. All instances of cis-mediation of trans-eQTLs with a mediation *p*-value <0.1 (16,648 trios) are listed in Supplemental Table S2, including trios containing transcripts with low mappability.

**Table 2.**
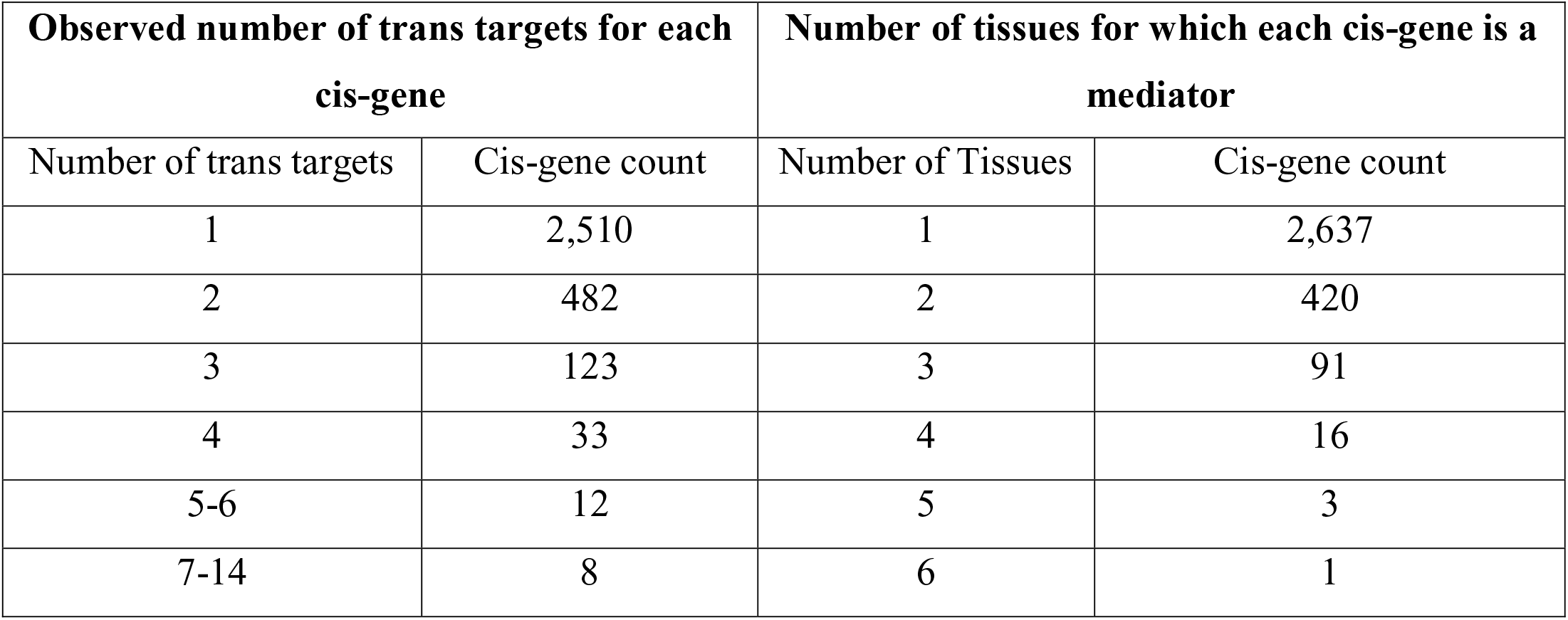
Frequency of cis-genes that mediate the effect of a trans-eQTL on multiple trans-genes or in multiple tissue types.

### Examples of mediation across tissues

In analyses restricting to cis and trans genes with mappability scores >0.95, one biologically interpretable example of a cis-gene that appears to mediate the effects of a trans-eSNPs in multiple tissues is the *IFI44L* gene on Chromosome 1 (Figure 4A). *IFI44L* is a cis-eGene in two GTEx tissues (cerebellar hemisphere and tibial nerve), and the cis-eSNPs associated with *IFI44L* expression are also associated with expression of multiple genes in trans in both cerebellar and tibial nerve tissue. *OAS1* is a trans target of these SNPs in both tissues, while other trans targets are observed in only cerebellar (*AGRN* and *PARP12*) or tibial nerve (*RSAD2, OAS2,* and *EPSTI1*). Below the mappability threshold of 0.95, we observe an additional potential trans targets of *IFI44L*, present in both cerebellar and tibial nerve tissue, *IFIT3* (mappability = 0.87). These relationships are depicted in Figure 4A.

**Figure 4.**
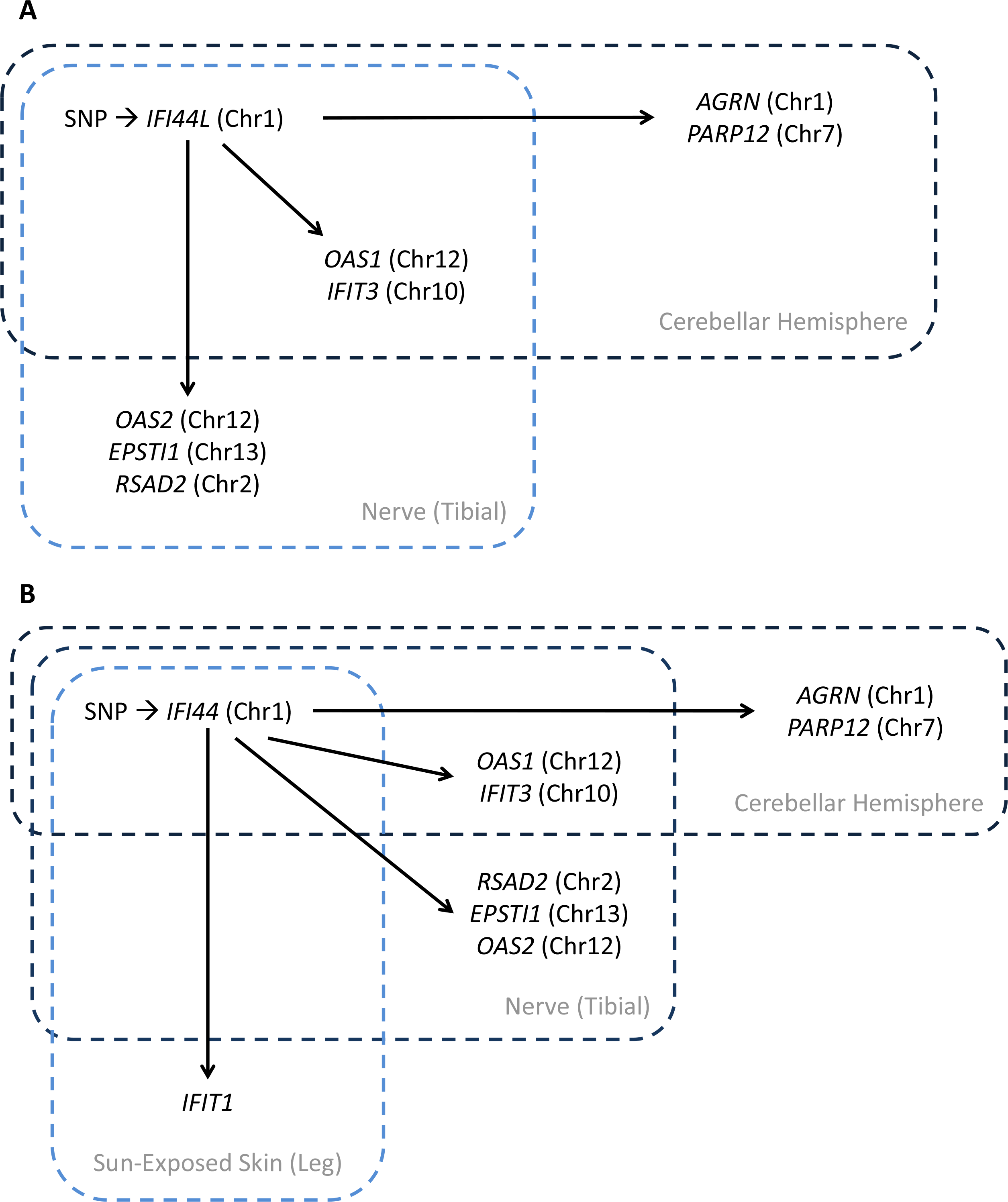
A biologically interpretable example of a cis-eGene (*IFI44L*) that appears to mediate the effects of trans-eSNPs in multiple tissues. The gene *IFI44L* (Panel A) resides <5 kb away from *IFI44* (B), and expression of these genes is associated with a common cis-eQTL that also impacts the expression of multiple genes in trans in multiple tissues. Both *IFI44* and *IFI44L* show statistical evidence of mediation for a similar set of interferon-related genes. Thus, based on this evidence, we infer that at least one of these genes is a cis-mediator, although cannot know which is (or if both are) the true mediator.

Interestingly, if we expand our analysis to include cis and trans genes with mappability >0.90, we detect *IFI44* (mappability of 0.93) as a cis-mediator regulating a nearly identical similar set of trans genes across three tissues: cerebellar hemisphere (*OAS1*, *IFIT2*, *AGRN*, and *PARP12*), tibial nerve (*OAS1, IFIT3, RSAD2*, and *EPSTI1*), and sun-exposed skin (*IFIT1*) (Figure 4B). *IFI44* and *IFI44L* are paralogs and reside adjacent to each other on 1p31.1. These genes are regulated by the same SNP in each tissue. It is highly unlikely that sequence similarity between these two genes causes our RNAseq-based expression measurements for *IFI44* (and/or *IFI44L*) to reflect the expression variation of both genes. The two regions of similarity shared between these two transcripts are 753 bp and 202 bp in length, and these regions share 69% and 67% similarity, respectively (Supplemental Fig S3). These two regions contain no identical sequences longer than ∼10 bp, making it impossible for an RNA-seq read (76 bp) to be ambiguous in terms of its mapping to *IFI44* vs. *IFI44L*. Furthermore, a prior study of array-based expression measures in endometrial cancer tissue reported genetic co-regulation of *IFI44* and *IFI44L* (in trans) (Kompass and Witte 2011).

Based on these observations, it is unclear which of these two genes is truly a cis-mediator of the observed trans-eQTLs (or if both are mediators). The causal cis-eSNP for *IFI44* (and/or *IFI44L*) appears to be different in different tissues, as the LD between the lead cis-eSNPs in cerebellar (rs12129932) and the lead eSNP in tibial nerve (rs74998911) is quite low with r^2^<0.01 in EUR 1000 Genomes data (Auton et al. 2015).

Regardless of the uncertainty whether *IFI44L* or *IFI44* (or both) is the true cis-hub of this trans-eQTL, nearly all of the genes involved in the putative regulatory pathways identified here are interferon-regulated/inducible genes, namely *OAS1, OAS2, IFIT1, IFIT3, IFI44, IFI44L, RSAD2*, and *AGRN* (Cheon and Stark 2009; Kyogoku et al. 2013). These genes have been previously reported to be co-expressed and/or co-regulated in various human cell types, including interferon-exposed fibroblasts and mammary epithelial cell lines (Cheon and Stark 2009), virus-infected airway epithelial cells cultures (Ioannidis et al. 2012), peripheral blood of individuals with acute respiratory infections (Zaas et al. 2009) as well as in both normal and cancerous human tissue (Cancer Cell Metabolism Gene DB, https://bioinfo.uth.edu/ccmGDB/). This previously-reported co-expression findings also extend to *EPSTI1* (Cheon and Stark 2009), the one gene we find to be a trans-target of *IFI44L* (and/or *IFI44*) that does not have a well-established function in immune response, providing additional evidence of an immune-related function for this gene.

Variation in the *IFI44L* gene is associated with risk for MMR (measles, mumps, and rubella) vaccination-related febrile seizures, with a missense variant in *IFI44L* showing the strongest association (Feenstra et al. 2014). Variation in *IFI44L* has also been implicated in schizophrenia risk (Ruderfer et al. 2014) as well as bipolar disorder (Chen et al. 2013). These findings suggest that the putative cross-tissue cis-hub identified here may be relevant to multiple neurological and psychological disorders, particularly those with etiologies related to immune function.

### Comparison of GMAC with other methods using simulated data

We evaluate the performance of the proposed GMAC in various simulated data scenarios. For each scenario described below, we simulated 1000 mediation trios (*L_i_*, *C_i_*, *T_j_*) for a sample size of n = 350, similar to the sample size of the GTEx data. Each mediation trio consists of a gene transcript *i* (*C_i_*), its cis-associated genetic locus (*L_i_*), and a gene transcript *j* (*T_j_*) in trans-association with the locus. Note that in the mediation analysis in this work (simulations and real data analysis), we consider only the trios with evidence of cis and trans-associations, *L_i_* → *ci* and *L_i_* → *T_j_*. We are interested in testing whether an observed trans-eQTL association is mediated by the cis-gene transcript, i.e., *L_i_* → *c_i_* → *T_j_*. We compared GMAC with other methods in different scenarios, including in the presence of confounders, common child variables, and intermediate variables. A common child variable is a variable that is affected by both *C_i_* and *T_j_* (Figure 2C). An intermediate variable is a variable that is affected by *C_i_* and affecting *T_j_*, that is, at least partially mediating the effects from *C_i_* to *T_j_* (Figure 2D).

#### Scenario 1: Under the null in the presence of common child variables

This is a scenario in which there is one common child variable for each pair of cis and trans gene transcripts (Figure 2C). In this scenario, adjusting for common child variables in mediation analyses would “marry” *C_i_* and *T_j_* and make *C_i_* appearing to be regulating *T_j_* even if there is no such effect (i.e., “collider bias”) (Greenland 2003) increasing the false positive rate for detecting mediation. Therefore, we consider it as “improper” to adjust for common child variable. We simulated a pool of independent and normally distributed variables **H**, with dimensionality being the same as the sample size of 350. For each of the 1000 mediation trios, we simulated the genetic locus *L_i_* under Hardy-Weinberg Equilibrium with a minor allele frequency of 0.1. Given *L_i_*, the cis-gene transcript *C_i_* and the trans-gene transcript *T_j_* are generated according to the models: *c_i_* = *β_i_*_0_*_c_* + *β_i_*_l_*_c_ L_i_* + *ϵ_ic_* and *T_j_* = *β_i_*_0_*_t_* + *β_i_*_l_*_t_ L_i_* + *ϵ_it_*. In this scenario, the trans-effect is not mediated by the cis-gene transcript. We let the parameters in the above models vary across the 1000 trios with *β_i_*_l_*_c_* sampled uniformly from 0.5 to 1.5, and the rest sampled uniformly from 0.5 to 1.0. The error terms *ϵ_ic_* and *ϵ_it_* are normally distributed. For each mediation trio, one candidate variable in **H** is randomly chosen to be the common child variable, *Z_j_*, and the effects of cis- and trans- gene transcripts on *Z_j_* are sampled uniformly from 1 to 1.5.

#### Scenario 2: Under the null in the presence of confounders

Scenario 2 is generated under the null in the presence of confounders (Figure 2B). Each candidate confounding variable has a 5% probability of being a true confounder of the cis-trans relationship for a randomly chosen proportion of trios where the proportion follows a uniform distribution from 0 to 0.2. This specification results in on average 1.85 confounders for each trio in our simulated data. Suppose for the i^th^ trio there are *n_i_* number of variables in **H** selected to be confounders, we denote the confounders as 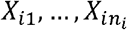. The cis-gene transcript *C_i_* and trans-gene transcript *T_j_* are generated according to the regression models 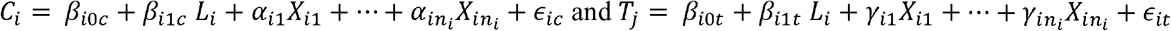. We let the parameters in the above models vary across the 1000 trios with similar parameter specification as before. In this scenario, there are no cis- to trans-gene mediation effects. Failure to adjust for confounders may induce false positive results, and it is improper to not adjust for confounders.

#### Scenario 3: Under the alternative in the presence of intermediate variables

We consider another scenario in which there is one intermediate variable for each cis-trans relationship (Figure 2D). For each mediation trio, we simulated the genetic locus and the cis-gene transcript as before and further simulated a child variable, *W_i_* of the cis-gene transcript. The trans-gene transcript, *T_j_* is then simulated to be affected by *W_i_*, according to 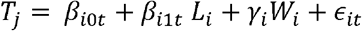. Note that the trans-gene *T_j_* is simulated to be affected by the cis-gene *C_i_* only via the intermediate variable *W_i_*. The mediation effects from cis to trans gene transcript (via *W_i_*) is non-zero in this scenario. Because *W_i_* is on the causal pathway from *C_i_* to *T_j_*, it is improper to adjust for *W_i_* and the adjustment will reduce or eliminate power to detect true mediation.

#### Scenario 4: Under the alternative in the presence of confounders

To compare with the existing approach that adjusts for a universal set of variables, we consider a scenario in which the dimensionality of potential confounding variables **H** is 100. For each trio, up to five variables in **H** are randomly selected to confound the cis trans gene relationship. The absolute effects of confounders on cis transcripts are sampled uniformly from 0.15 to 0.5 with a 50% probability to be negative. And the effects of confounders on trans transcripts are sampled uniformly from 0.15 to 0.5 with all to be positive. We set the effect of cis transcript on trans transcript to be 0.1, i.e., non-zero mediation effects. When the number of potential confounding variables is large, although one may still adjust them all for the simulated sample size, this adjustment is inefficient and may hurt the power.

#### Simulation Results

For each scenario, we compared the results based on the following methods: 1) Oracle adjustment, which correctly adjusts for the true confounders but no child or intermediate variables in the mediation test; 2) The GMAC algorithm; and 3) Improper (or inefficient) adjustment, which corresponds to incorrectly adjusting for the common child variables in scenario 1, failure to adjust for confounders in scenario 2, incorrectly adjusting for the intermediate variables in scenario 3, and universally adjusting for all variables in **H** in scenario 4, including variables which are not true confounders. Table 3A and 3B show the true type I error rates at the significance levels of 0.01 and 0.05 in scenario 1 and 2 respectively. As expected, adjusting for child variables “marries” the cis- and trans-genes in the mediation test, resulting in inflated rates of false positive findings. Failure to adjust for confounding also leads to inflated type I error rates. In contrast, both the oracle and GMAC adjustment control the type I error rates. Table 3C shows that when the power to detect mediation is high (by Oracle and GMAC), incorrectly adjusting for an intermediate variable in this setting greatly reduces the power to detect mediation. In comparison, GMAC correctly filters out most of the true intermediate variables in the mediation tests and maintains power comparable to oracle adjustment. Table 3D shows that GMAC has comparable power than oracle adjustment in scenario 4. In our simulation, 2,962 out of 3,023 generated confounders across the 1000 mediation trios are correctly selected. In comparison, adjusting for all variables in the pool of confounders is inefficient and reduces power to detect mediation.

**Table. 3.**
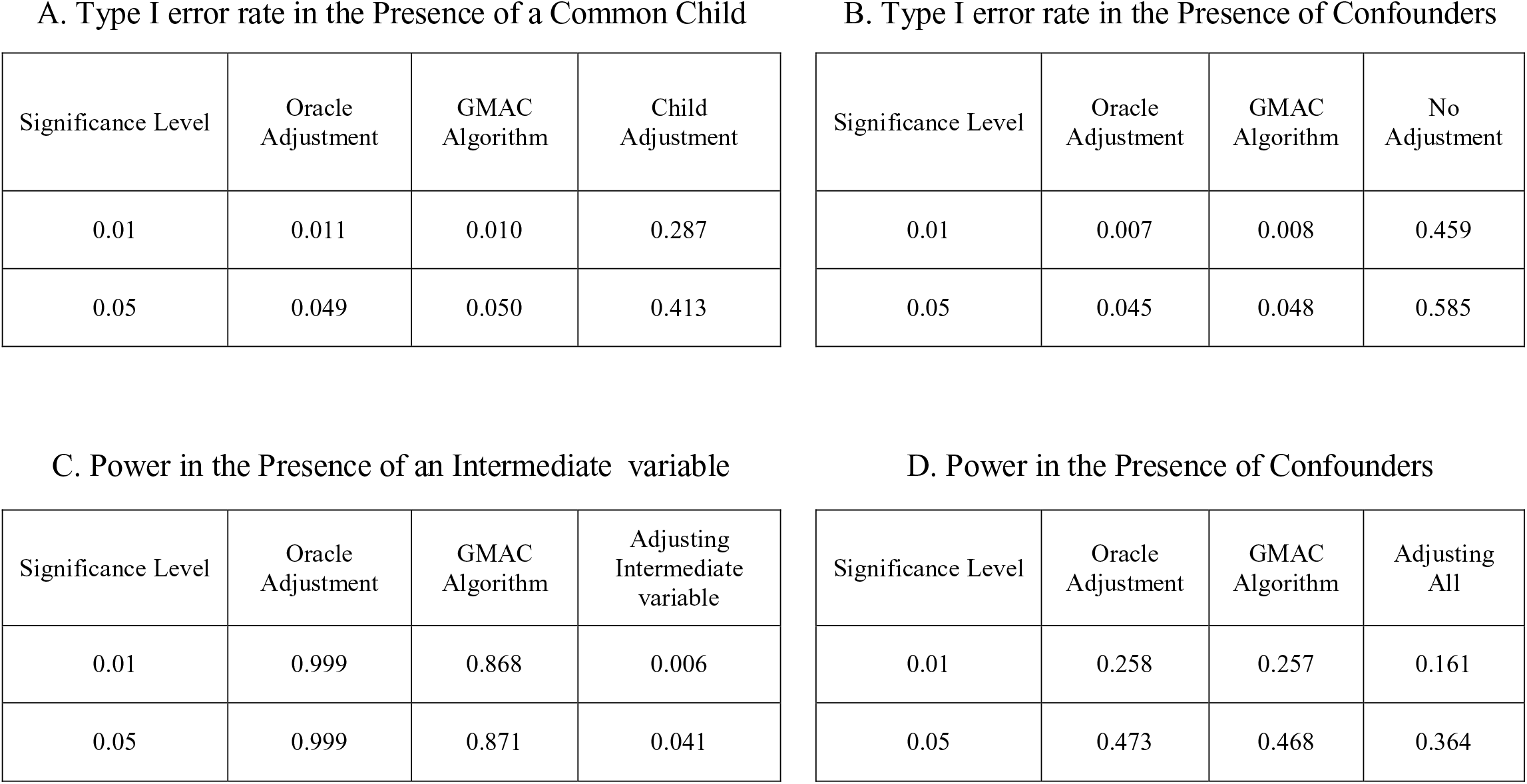
Comparison of the type I error rate and power of GMAC compared to other methods for mediation analysis under the null (A and B) and the alternative (C and D) hypotheses, based on simulated data.

## DISCUSSION

In this work, we have developed the GMAC algorithm for conducting mediation analysis to identify cis- transcripts that mediate the effects of trans-eQTLs on distant genes. We address a central problem in mediation analysis, “mediator-outcome confounding,” by developing an algorithm that can 1) search a very large pool of variables (surrogate and/or measured) for variables likely to have confounding effects and 2) adaptively adjust for such variables in each mediation test conducted. We acknowledge that we cannot make definitive causal claims regarding any of the mediation/regulatory relationships for which we detect evidence. Instead, the focus of our work is to strengthen the evidence that mediation analysis can provide, and to identify candidate cis-hub genes likely to mediate the effects of eQTLs on many trans- genes within or across human tissue types. Cis-hub genes are likely to be key players in the regulatory networks relevant to human disease, thus it is important that we understand their patterns of regulation. By applying GMAC to 44 human tissues from the GTEx project, we are able to characterize cis-hubs with potential disease relevance by aggregating information across many different tissue types. Analyses of simulated data show that the GMAC algorithm improves the power to detect true mediation compared with existing methods, while controlling the true false discovery rate.

In analyses of GTEx data, over 20% of cis-mediators we observe appear to mediate the effects of a trans-eQTL on multiple genes, but the vast majority of these cis-hubs are either tissue-specific (i.e., mediating multiple trans-genes in a single tissue type) or have unique trans targets in each tissue type. We provided one example of a biologically plausible multi-tissue cis-hub, whereby a cis-mediator of a trans-eQTLs appears to have common trans targets across multiple tissue types. The cis-hub identified (*IFI44L*) has potential relevance for neurological and psychological disorders, particularly those with etiologies related to immune function, demonstrating the potential value of our approach for understanding disease-relevant pathways.

One innovative aspect of this work is our algorithm that rigorously addresses the problem of “mediator-outcome confounding” in the context of genomic mediation analysis. In eQTL-based mediation analysis, potential confounders of the cis-trans association include demographic and environmental factors, as well as a wide array of biological phenomenon, such as expression of specific genes or other biological processes that may be represented by the expression of sets of genes. Neglecting to control for such confounding variables can lead to substantial bias in estimates of mediation, resulting in spurious findings, as we have described previously (Pierce et al. 2014). Considering the complexity of the biological systems under study, as well as the diversity of the GTEx donors, a careful control for such confounding variables is critically important.

Most existing methods control for confounding variables by constructing a set of variables that represent the largest components of variation in the transcriptome and adjusting for the selected set for all tests conducted. However, only when a variable is causally related to both the cis- and trans-genes (as shown in Figure 2B) will the variable potentially show confounding effects in mediation analysis. GMAC adaptively selects a set of confounding variables for each trio undergoing mediation analysis, enabling large-scale genomic mediation analyses adjusting only for the confounding variables that could potentially bias a specific mediation estimate. As opposed to adjusting for all known covariates, our strategy of selecting only potential confounders for adjustment purposes is important for three reasons: 1) Adjusting for fewer variables increases power (i.e., fewer degrees of freedom; see Supplemental Table S1); 2) The number of variables from which one selects covariates could be extremely large (e.g., all expressed genes), making adjustment for all covariates impossible; and 3) inadvertently adjusting for “common child” or intermediate variables can result in substantial biases. In this work we select potential confounders from all expression PCs, but one could also select from among transcripts that are not well-represented by PCs. By efficiently selecting confounders from a very large pool of potential variables, GMAC improves both power and precision in mediation analyses.

There are several limitations of our approach and its application to GTEx data. First, when working with real genomic data, we can never be sure that we have measured and accounted for all possible mediator-outcome confounding. Potential confounders include participant characteristics, environmental factors, tissue micro-environmental factors, as well as a wide array of biological factors which may or may not be captured by the expression data being analyzed. Second, in the analysis presented here, we only consider the trios with both strong cis- and trans-eQTL effects. For any given tissue type we are analyzing, our sample size is too small for robust genome-wide detection/analysis of trans-eQTLs. As such, the mediation trios we considered are only a subset of the true mediation trios in the genome. And the small sample sizes may also result in underpowered mediation tests. As the sample size of GTEx increases, future studies will have increased power to identify cis-mediators using GMAC. Third, we did not consider the full complexity of gene isoforms and splice variants in this work; future studies should consider the possibility of mediation relationships that are isoform-specific. Lastly, some trans-eQTLs may not be mediated by variation in the expression of a cis-gene. Other potential mediating mechanism could include variation in coding sequence, physical inter-chromosomal interaction, or variation in non-coding RNA. Our work is not intended to identify and analyze such trans-eQTLs, as we perform trans-eQTL analyses using only SNPs known to be cis-eSNPs.

It is important to note that our expectation is that most trans-eQTLs are fully-mediated by a transcript that is regulated in cis by the causal trans-eQTL variant. We did not observe “complete mediation” (i.e., % mediation = 100%) for the majority of the significant mediation *p*-values we observed. However, as we have explained and demonstrated previously (Pierce et al. 2014), full mediation will be observed as partial mediation in the presence of mediator measurement error and/or imperfect LD between the causal variant and the variant used for analysis purposes. Thus, considering RNA quantification is not error free and causal variants are often unknown, we expect to often observe partial mediation when full mediation is present.

We also demonstrate that it is critical to consider mappability for both cis and trans genes involved in mediation analysis. For genes containing sequences that do not uniquely map to the human transcriptome, it is possible that gene expression measures may be comprised of signals coming from multiple genes, which can produce false positives in mediation analysis, including spurious detection of cis-hubs and cross-tissue cis-mediators.

Our application of the GMAC algorithm to the multi-tissue expression data from GTEx provides a unique cross-tissue perspective on cis-mediation of trans-regulatory relationships across human tissues. This multi-tissue perspective is important because observing mediation relationships that are consistent across multiple tissues provides confidence that a significant mediation *p*-value reflects a true instance of mediation. For the “cis-hub” genes and genes that appear to be cis-mediators in multiple tissues, further investigation is warranted, as these genes may have many regulatory relationships that we are not powered to detect in this work. Thus, a multi-tissue mediation analysis approach has the potential to increase power to identify true mediators while controlling for false positives. In future work, attempts at joint analyses of multiple tissue types may provide a more complete picture of the cross-tissue and tissue-specific trans-regulatory mechanisms. The GMAC approach described here will be a valuable tool for such studies, as well any future studies that aims to understand the relationships among cis- and trans-eQTLs and characterize the biological mechanisms and networks involved in human disease biology.

We have developed an R GMAC package to perform the proposed genomic mediation analysis with adaptive selection of confounding variables. The package is currently available through R CRAN.

## METHODS

### Bio-specimen collection and processing of GTEx data

A total of 7,051 tissues samples were obtained from 44 distinct tissue types from 449 post-mortem tissue donors (with 65.6% male). Those donors were from multiple ethnicity groups, spanned a wide age range, and have various reasons of death (see GTEx portal for descriptive statistics). Donor enrollment, consent processes, biospecimen collection and processing have been described previously (Lonsdale et al. 2013; Ardlie et al. 2015). Briefly, each tissue sample was preserved in PAXgene tissue kit and the stored as both frozen and paraffin embedded tissue. Total RNA was isolated from PAXgene fixed tissues samples using the PAXgene Tissue mRNA kit. For whole blood, Total RNA was isolated from samples collected and preserved in PAXgene blood RNA tubes.

Blood samples were used as the primary source of DNA. Genotyping was conducted using the Illumina Human Omni5-Quad and Infinium ExomeChip arrays. Standard QC procedures were performed using the PLINK software (Purcell et al. 2007) and genotype imputation was performed using the IMPUTE2 software (Howie et al. 2009) and reference haplotypes from the 1000 Genomes Project (Auton et al. 2015). The first three PCs representing ancestry (Price et al. 2006) were included as covariates in all analyses.

RNA-seq data was generated for RNA samples with a RIN value of 6 or greater. Non-strand specific RNA sequencing was performed using an automated version of the Illumina TruSeq RNA sample preparation protocol. Sequencing was done on an Illumina HiSeq 2000, to a median depth of 78M 76 bp paired-end reads per sample. RNA-seq data was aligned to the human genome using TopHat (Trapnell et al. 2009). Gene-level expression was estimated in RPKM units using RNA-SeQC (DeLuca et al. 2012). RNA-seq expression samples that passed various quality control measures (as previously described) were included in the final analysis dataset.

### Mappability of transcripts

Because non-uniquely mapping reads can result in false positive eQTLs, we use the mappability of each gene as a quality control filter, as described in Jo et al (the GTEx “trans paper”). The mappability was calculated as follows: Mappability of all k-mers in the reference human genome (hg19) computed by ENCODE (The ENCODE Project Consortium 2012) was downloaded from the UCSC Genome Browser (accession: wgEncodeEH000318, wgEncodeEH00032) (Rosenbloom et al. 2013). The exon- and UTR- mappability of a gene was computed as the average mappability of all k-mers in exons and UTRs, respectively. We used k = 75 for exonic regions, as it is the closest to GTEx read length among all possible k’s. UTRs are generally quite small, so k = 36 was used, the smallest among all possible k’s. Mappability of a gene was computed as the weighted average of its exon-mappability and UTR-mappability, with the weights being proportional to the total length of exonic regions and UTRs, respectively.

### The selection of trios for mediation tests

In the mediation analysis presented in this work, we consider only the trios with evidence of cis and trans-associations, *L_i_* → *C_i_* and *L_i_* → *T_j_*. The identification of cis-eQTLs is described elsewhere (cite Aguet et al. 2016, GTEx companion paper, unpublished) (Ardlie et al. 2015). For genes with multiple cis-eSNPs as eQTLs, only one cis-eSNP for each gene (i.e., the high-quality SNP with the smallest P-value) was selected and was included in the subsequent trans-eQTL and mediation analyses. The complete cis-eQTL list is available through GTEx portal (https://gtexportal.org/) and all data can be obtained through dbGaP (phs000424.v6.p1).

Furthermore, using Matrix eQTL (Shabalin 2012) we conducted genome-wide trans-eQTL analyses restricting to the cis-eSNPs described above and examining association for all genes located at least 1Mb away from the cis-eSNPs. Up to 35 PEER factors and other covariates were adjusted. In each tissue type, when a trans-association *p*-value is less than 10^-5^, the eSNP, its corresponding cis-transcript, and the trans-transcript were treated as a candidate trio and were retained for mediation analysis.

### The GMAC algorithm

In order to identify cis-mediators of trans-eQTLs across the genome, we propose the GMAC algorithm (Figure 2A). Here we present a brief description of each step. A detailed description and justification for each can be found in the Supplemental Methods. Specifically,

- <u>Step 0</u>. We focus on only the trios (*L_i_*, *C_i_*, *T_j_*) in the genome showing both cis- and trans-eQTL associations, i.e., *L_i_ → C_i_* and *L_i_* → *T_j_*. Consider a pool of candidate variables **H** consisting of either real covariates, constructed surrogate variables, or both.
- <u>Step 1. Filter out common child and intermediate variables from the pool of potential confounders</u>. For each trio (*L_i_*, *C_i_*, *T_j_*) we calculate the marginal associations of variables in **H** to *L_i_* and filter the ones with significant associations at the 10% FDR level. As shown in Figure 2B-D, common child and intermediate variables are directly associated with *L_i_*, while confounders are assumed to be unassociated with *L_i_*. Note that since genetic loci are “Mendelian randomized” (Smith and Ebrahim 2003), without loss of generality we assume the confounders are not associated with *L_i_*. Let **H_ij_** denote the retained pool of candidate variables specific to the trio (*L_i_*, *C_i_*, *T_j_*).
- <u>Step 2. Adaptively select confounders.</u> For each trio and each of its potential confounding variables in **H_ij_**, we calculate the *p*-value of the overall F-test to assess the association of the variable to at least one of the cis- and trans-transcripts. Considering the *p*-values for one potential confounding variable to all trios as one stratum, we apply a 10% FDR significance threshold to each stratum (each column in Figure 2E) – a stratified FDR approach (Sun et al. 2006). The significant variables corresponding to a trio (each row in Figure 2E) will be selected in the mediation analyses as the adaptively selected confounders specific to that trio (see Supplemental Methods for details). Let **X_ij_** denote the list of adaptively selected confounding variables for the trio, (*L_i_*, *C_i_*, *T_j_*).
- <u>Step 3.Test for mediation.</u> For each trio and its adaptively selected confounder set, we calculate the mediation statistic as the Wald statistic for testing the indirect mediation effect *H_0_: β_1_ = 0* based on the following regression regressing the trans-gene expression levels on the cis-gene expression levels adjusting for the cis-eQTL, other covariates and selected confounders:

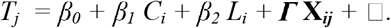 We perform within-genotype group permutation on the cis-gene transcript at least 10,000 times and re-calculate each null mediation statistic based on the locus, a permuted cis-gene transcript, and the trans-gene transcript, (*L_i_*, *C_i0_*, *T_j_*). Figure 2F shows the expression variation patterns of a hypothetical mediation relationship *L_i_ → C_i_ → T_j_* on the left panel, and a null relationship entailed by (*L_i_*, *C_i0_*, *T_j_*) with *L_i_ → C_i0_* and *L_i_ → T_j_* but no mediation. It justifies that by permuting the cis-gene expression levels within each genotype group, one maintains the cis-associations while breaks the potential mediation effects from the cis- to the trans-gene transcript (i.e., conditional correlations of cis and trans transcript conditioning on *L_i_*, or correlation within each genotype group). We calculate the *p*-value of mediation for the trio (*L_i_*, *C_i_*, *T_j_*) by comparing the observed mediation statistic with the null statistics.

The proposed algorithm is superior to existing approaches for mediation analysis that adjust a universal set of variables for all trios. GMAC avoids the adjustment of common child variables, intermediate variables and unrelated variables in genomic mediation analysis, and it is able to search a much larger pool of variables for potential confounders, not just those captured by the top few surrogate variables or PCs.

## SOFTWARE AVAILABILITY

The R software package GMAC is available in the Supplemental Materials and also online through R CRAN https://cran.r-project.org/.

## ACKNOWLEDGMENTS

We would like to thank Alexis Battle for providing the estimates of mappability used in this work. The Genotype-Tissue Expression (GTEx) Project was supported by the Common Fund of the Office of the Director of the National Institutes of Health. Additional funds were provided by the NCI, NHGRI, NHLBI, NIDA, NIMH, and NINDS. Donors were enrolled at Biospecimen Source Sites funded by NCI\SAIC-Frederick, Inc. (SAIC-F) subcontracts to the National Disease Research Interchange (10XS170), Roswell Park Cancer Institute (10XS171), and Science Care, Inc. (X10S172). The Laboratory, Data Analysis, and Coordinating Center (LDACC) was funded through a contract (HHSN268201000029C) to The Broad Institute, Inc. Biorepository operations were funded through an SAIC-F subcontract to Van Andel Institute (10ST1035). Additional data repository and project management were provided by SAIC-F (HHSN261200800001E). The Brain Bank was supported by a supplements to University of Miami grants DA006227 & DA033684 and to contract N01MH000028. Statistical Methods development grants were made to the University of Geneva (MH090941 & MH101814), the University of Chicago (MH090951, MH090937, MH101820, MH101825), the University of North Carolina - Chapel Hill (MH090936 & MH101819), Harvard University (MH090948), Stanford University (MH101782), Washington University St Louis (MH101810), and the University of Pennsylvania (MH101822). The data used for the analyses described in this manuscript were obtained from: the GTEx Portal in January of 2015. This work was supported by National Institutes of Health grants R01 GM108711, U01 HG007601, and R01 MH101820.

## DISCLOSURE DECLARATION

The authors declare no competing financial interest.

